# Genome Graphs

**DOI:** 10.1101/101378

**Authors:** Adam M. Novak, Glenn Hickey, Erik Garrison, Sean Blum, Abram Connelly, Alexander Dilthey, Jordan Eizenga, M. A. Saleh Elmohamed, Sally Guthrie, André Kahles, Stephen Keenan, Jerome Kelleher, Deniz Kural, Heng Li, Michael F. Lin, Karen Miga, Nancy Ouyang, Goran Rakocevic, Maciek Smuga-Otto, Alexander Wait Zaranek, Richard Durbin, Gil McVean, David Haussler, Benedict Paten

## Abstract

There is increasing recognition that a single, monoploid reference genome is a poor universal reference structure for human genetics, because it represents only a tiny fraction of human variation. Adding this missing variation results in a structure that can be described as a mathematical graph: a genome graph. We demonstrate that, in comparison to the existing reference genome (GRCh38), genome graphs can substantially improve the fractions of reads that map uniquely and perfectly. Furthermore, we show that this fundamental simplification of read mapping transforms the variant calling problem from one in which many non-reference variants must be discovered de-novo to one in which the vast majority of variants are simply re-identified within the graph. Using standard benchmarks as well as a novel reference-free evaluation, we show that a simplistic variant calling procedure on a genome graph can already call variants at least as well as, and in many cases better than, a state-of-the-art method on the linear human reference genome. We anticipate that graph-based references will supplant linear references in humans and in other applications where cohorts of sequenced individuals are available.

## Introduction

The human reference genome, completed in draft form in 2001 and revised several times subsequently ^1,2^, is the single most important resource used in human genetics today. It acts as a universal coordinate system and as such is the space in which annotations (genes, promoters, etc.) and genetic variants are described ^3–5^. It is also the target for read mapping, and, downstream of this mapping, is used for functional assays and variant calling pipelines ^6,7^.

The contemporary definition of a reference genome is completely linear: a single monoploid assembly of the genome of a species. A key limitation of the linear human reference genome (the set of chromosome scaffolds) is that it is but a single genome. As such, it is an imperfect lens through which to study our population’s variation; there exist variants and annotations that can not be easily described with respect to the reference genome ^8,9^. Furthermore, as a target for mapping and interpretation it introduces a reference allele bias: a tendency to over-report alleles present in the reference genome and under-report other alleles ^10,11^. To mitigate these issues, recent versions of the reference genome assembly, such as GRCh38, have contained “alternate locus” sequences (“alts”): extra sequence representations of regions of the human genome considered to be highly polymorphic, anchored at their ends to locations within the “primary” (monoploid) reference assembly. Such a structure, which contains multiple partially-overlapping sequence paths, can be considered a form of mathematical graph. The explicit use of graphs in biological sequence analysis has a long history, notably for sequence alignment ^12^, sequence assembly ^13,14^, assembly representation (as in FASTG and now GFA)^15,16^, substring indexes (which are often thought of in terms of suffix trees or similar data structures) ^6,17^, and transcript splice graphs ^18^. Recently the notion of graphs for representing genomes has been considered explicitly ^12,19,20^, and work has been done towards using these graphs as references ^21^. The alternate loci currently used are just one way to extend the linear reference genome into a genome graph; many other ways are possible. In this work, conducted by a task team of the Global Alliance for Genomics and Health, we experiment with different methods for graph construction and testing the utility of different graphs for read mapping and variant calling. This work is the first study of its kind that we are aware of. We attempt to test the simple hypothesis that adding data into the reference structure—in effect, adding to the “reference prior” on variation extant in the population—will result in improved genome inferences.

## Results

There are many possible types of genome graph; here we use *sequence graphs*. The nodes of a sequence graph are a set of DNA sequences. Each node is therefore a string of nucleotide characters, called positions, giving the sequence of the node’s forward strand. We call the terminal 5’ and 3’ ends of this strand the *sides* of the node. Each edge in the graph is an unordered pair of sides, representing a (potential) bond between two sides of a pair of nodes. This is a bidirected graph representation, because features of the edge indicate to which side of a node (sequence), 5’ or 3’, each end of the edge connects (Fig. 1)^22^. Other representations of genome graphs, such as the directed acyclic representation, can be useful; see Supplementary Section 1. A longer DNA sequence can be represented as a *thread* within a sequence graph, beginning in one oriented node, ending in the same node or another, and in between walking from node to node, with the rule that if the walk enters a node on one side it exits through the other side.

**Figure 1:**
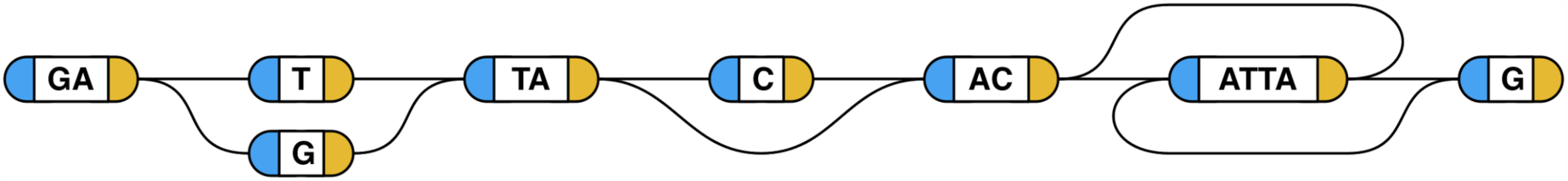
Example sequence graphs. Each node holds a string of bases. An edge can connect, at each of its ends, to a base on either the left (5’, blue) or the right (3’, yellow) side of the node. When reading through a thread to form a DNA sequence, a valid walk must leave each node via the opposite side from that through which it was entered; a node’s sequence is read as reverse-complemented if the node is entered on the 3’ side. One thread that this graph spells out (reading from the left side of the leftmost sequence to the right side of the rightmost sequence, along the nodes drawn in the middle) is the sequence “GATTACACATTAG”. Straying from this path, there are three variants available: a substitution of “G” for “T”, a deletion of a “C”, and an inversion of “ATTA”. If all of these detours are taken, the sequence produced is “GAGTAACTAATG”. All 8 possible threads from the leading G to the trailing G are allowed.

To evaluate the utility of sequence graphs we invited teams to construct and evaluate graphs for five test regions of the human genome: the major histocompatibility complex (MHC), the killer cell immunoglobulin-like receptors (LRC_KIR) region, the spinal muscular atrophy (SMA) locus, and the BRCA1 and BRCA2 genes. MHC, SMA and LRC_KIR are all regions with alternate loci represented in GRCh38, while BRCA1 and BRCA2 represent more typical human genes. Regions ranged from 81 kilobases in size with a single gene (BRCA1) to 5.0 megabases in size with 172 genes (MHC). We considered graphs from five teams built with eight different pipelines (Table 2). For each region we provided a set of long, high quality input sequences from which to construct graphs (Table 1), but also encouraged the creation of graphs using additional data of the builder’s choice. Some graphs were built based upon existing variant calls, such as the 1000 Genomes calls used to construct the 1 KG graph ^5^. Graphs were also built with a wide variety of different algorithmic approaches (Table 2). Three control graphs were constructed for each region as points of comparison. The Primary graphs contain just the single, linear reference path from GRCh38. The Unmerged graphs consist of just the set of provided sequences, each represented as a disjoint path. The Scrambled graphs (see Online Methods) are essentially identical topologically to the 1KG graphs, but with structures shifted to create false variants. These graphs acted as a negative control for the effects of adding nonsense variation to the graphs.

**Table 1:**
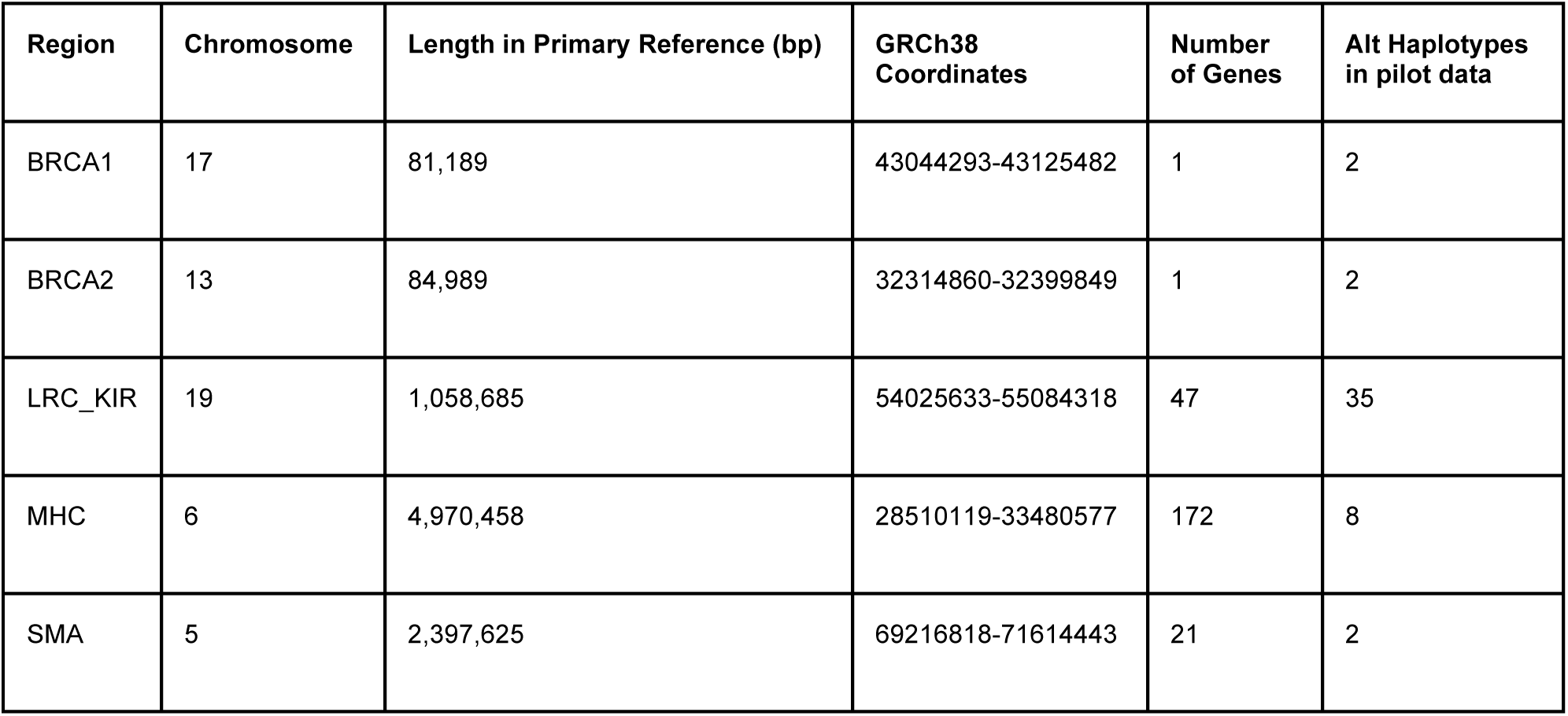
**Pilot Regions**. Selected test cases represent a sampling of both typical and challenging genomic regions.

| Region | Chromosome | Length in Primary Reference (bp) | GRCh38 Coordinates | Number of Genes | Alt Haplotypes in pilot data |
| --- | --- | --- | --- | --- | --- |
| BRCA1 | 17 | 81,189 | 43044293-43125482 | 1 | 2 |
| BRCA2 | 13 | 84,989 | 32314860-32399849 | 1 | 2 |
| LRC_KIR | 19 | 1,058,685 | 54025633-55084318 | 47 | 35 |
| MHC | 6 | 4,970,458 | 28510119-33480577 | 172 | 8 |
| SMA | 5 | 2,397,625 | 69216818-71614443 | 21 | 2 |

**Table 2:** **Genome Graph Submissions**. Submissions were collected from a variety of institutions, and showcase a variety of graph construction methods.

| Submissions using pilot data |  |  |  |
| --- | --- | --- | --- |
| Submission | Team | Short Name | Description of Algorithm |
| Cactus | UCSC | Cactus | Graph-based multiple sequence aligner <sup>23</sup>. |
| Camel | UCSC | Camel | Creates graphs progressively by mapping using context schemes <sup>24</sup>. |
| De Bruijn Graph (k=63) | MSKCC | De Bruijn 63 | Forms a De Bruijn graph of input data with k=63, then converts to a sequence graph. |
| Population Reference Graph | Oxford | PRG | Creates a graph from a K-mer-based HMM description of a region <sup>19</sup>. |
| Seven Bridges | Seven Bridges | 7BG | Multiple genome alignment. |
| Submissions using other data |  |  |  |
| Submission | Team | Short Name | Description of Algorithm |
| 1000 Genomes SNP Graph | Sanger/UCSC | 1KG | Generated using vg construct on a VCF containing variants identified in the 1000 genomes project. Platinum genome samples were not included, to avoid circularity in variant evaluation. |
| 1000 Genomes Haplotype 50 | Sanger/UCSC | 1KG Haplo 50 | Adapted form of 1KG graph in which phasing information is used to reduce the number of unobserved recombinations represented by the graph. 50 is the number of bases two variants can be apart to be considered for this phasing. |
| Scrambled 1000 Genomes | Sanger/UCSC | Scrambled | Generated by shifting all the variants in the standard 1KG graph 200 bp downstream. |

## Graph Read Mapping

To evaluate the utility of sequence graphs for read mapping we used the software program vg^25^, which contains a mapping algorithm capable of mapping to a fully general and potentially cyclic sequence graph (see Supplementary Section 2). We mapped all relevant reads (see Online Methods) from 1000 Genomes Phase 3 low coverage samples to each graph. We found that vg was able to map almost all such reads to the graphs (Supplementary Fig. 3).

Relative to the primary graph, a graph containing more of the variants should produce an increase in the fraction of reads that map perfectly (without either substitutions or indels) to at least one place. For BRCA2 we see a relative increase of 7.3% in the median number of reads mapping perfectly to the 1 KG graph vs. the Primary graph, but for MHC the increase is 20% (Fig. 2 top row, Supplementary Section 3, Supplementary Fig. 1). The increase for BRCA2 is close to what would be expected if the sequence graph contained the majority of polymorphisms for a typical region of the genome (Supplementary Section 3), while the larger increase for MHC is likely due to a greater degree of polymorphism ^11^. Similar, slightly smaller increases are seen for graphs built from other, smaller collections of variants. The scrambled graphs do not show significant gains, thus indicating that the effect is specific to graphs containing known variation. Furthermore, the overall substitution rate between reads and the experimental graphs was observed to decrease, relative to the rate between the reads and the Primary control graph. In the highest-performing graphs the decline is slightly below the bounds of previous read substitution error rate estimates of 0.7-1.6% ^5,26–28^ (Fig. 2 second row; see Supplementary Section 4 and Supplementary Fig. 4). The decrease in indel rate (Fig. 2 third row) moving from the Primary graph to the 1KG graph is comparable to estimates of the human indel polymorphism rate^29^ (Supplementary Section 5).

**Figure 2:**
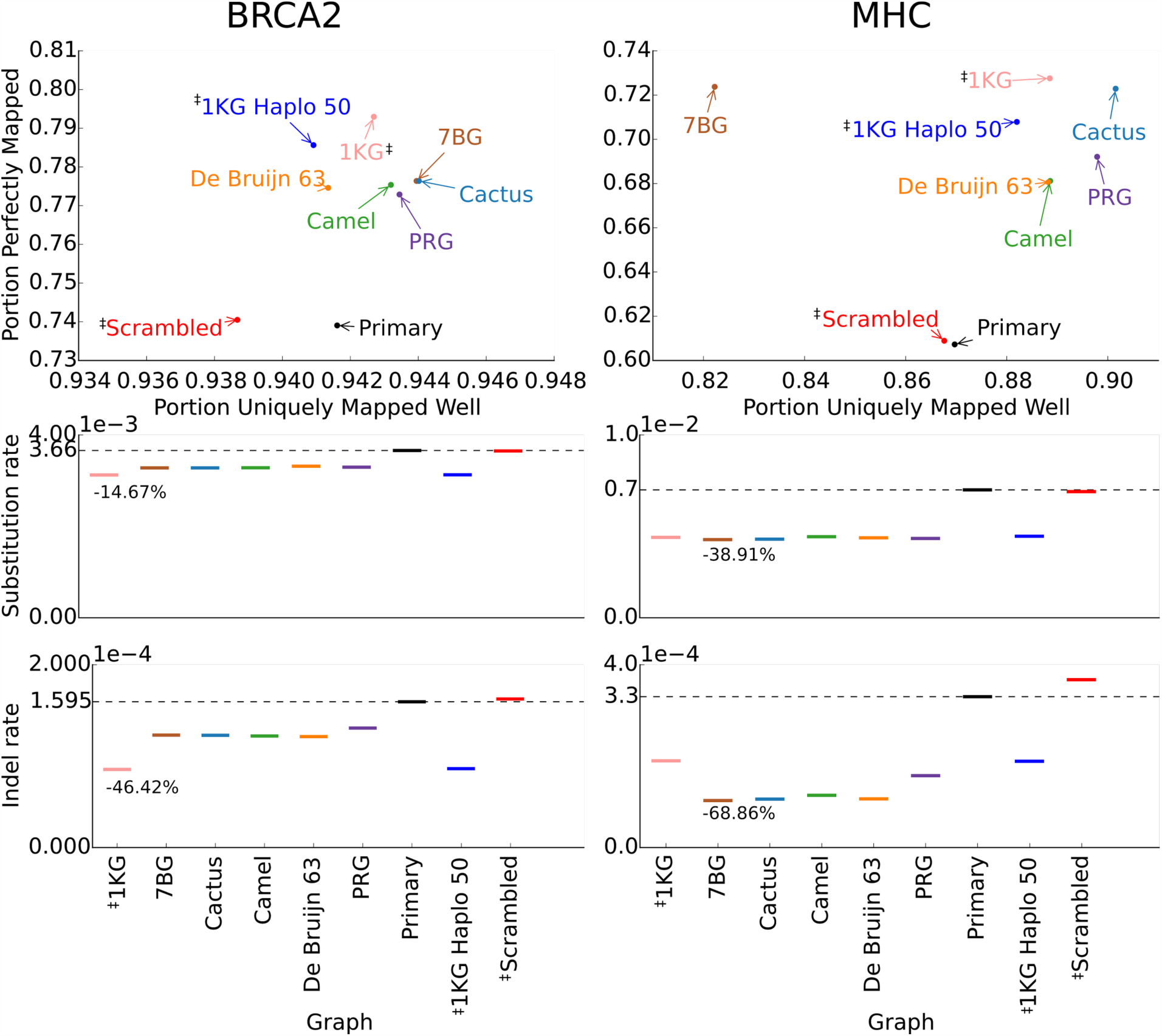
Mapping reads to sequence graphs. Results for the 1000 Genomes Phase 3 low coverage samples against the BRCA2 and MHC graphs. The median per-sample portion of reads that are mapped perfectly (Y axis), and the median per-sample portion of reads that are mapped with a unique, obviously-best alignment (X axis) are both visible in the top row. The median per-sample substitution rate for a primary mapping, computed per aligned base, is shown in the second row. The median per sample frequency of indels in primary mapped reads, computed per read base, is given in the third row. The horizontal black line represents the result for the primary reference graph in the region. The ‡ symbol marks graphs generated using additional data beyond the provided reference and alternate sequences. The unmerged graphs are excluded because very few reads mapped uniquely to them.

The median fraction of reads that uniquely map increases for many of the graphs, relative to the primary and scrambled graphs. For example, in the Cactus graph, an increase of 0.26% is observed in BRCA2, and an increase of 3.7% is observed in the MHC. No such increase in unique mapping is seen for the comparably complex scrambled graph. Unique mapping is defined as having a good primary mapping and no reasonable secondary mapping (see Supplementary Section 3 and Supplementary Fig. 2).

To test if the choice of sequence graph reference affected population level reference allele bias, we binned samples by 1KG super-population and looked at the difference in perfect mapping between the 1KG graph and the Primary graph for each subpopulation. We find a small but significant difference in perfect mapping increase between super-populations for most regions (Supplementary Section 6, Supplementary Fig. 5), but we also find relatively large differences in absolute rates of perfect mapping (Supplementary Fig. 6). These latter differences suggest that super-population may be confounded with sequencing batch, making drawing conclusions from this analysis quite difficult.

## Graph Variant Calling

We implemented a comprehensive, albeit basic, variant calling pipeline based on the Samtools pileup approach ^30^, modified to work with sequence graphs (see Online Methods for more details). In summary (Fig. 3), reads are mapped to the reference graph or *base graph* and a pileup is computed at each graph position and edge. An *augmented graph* is created by extending the base graph with additional sequences and edges representing possible variants determined from the pileups. This graph is then analyzed for *ultrabubbles* (acyclic, tip-free subgraphs connected to the rest of the graph by at most 2 nodes) which are treated as sites for genotyping^31^. Finally, thresholding heuristics are used to emit a set of genotypes with sufficient read support, one for each site, expressed in the coordinates of the GRCh38 primary reference path as embedded in the graph (see Online Methods).

**Figure 3:**
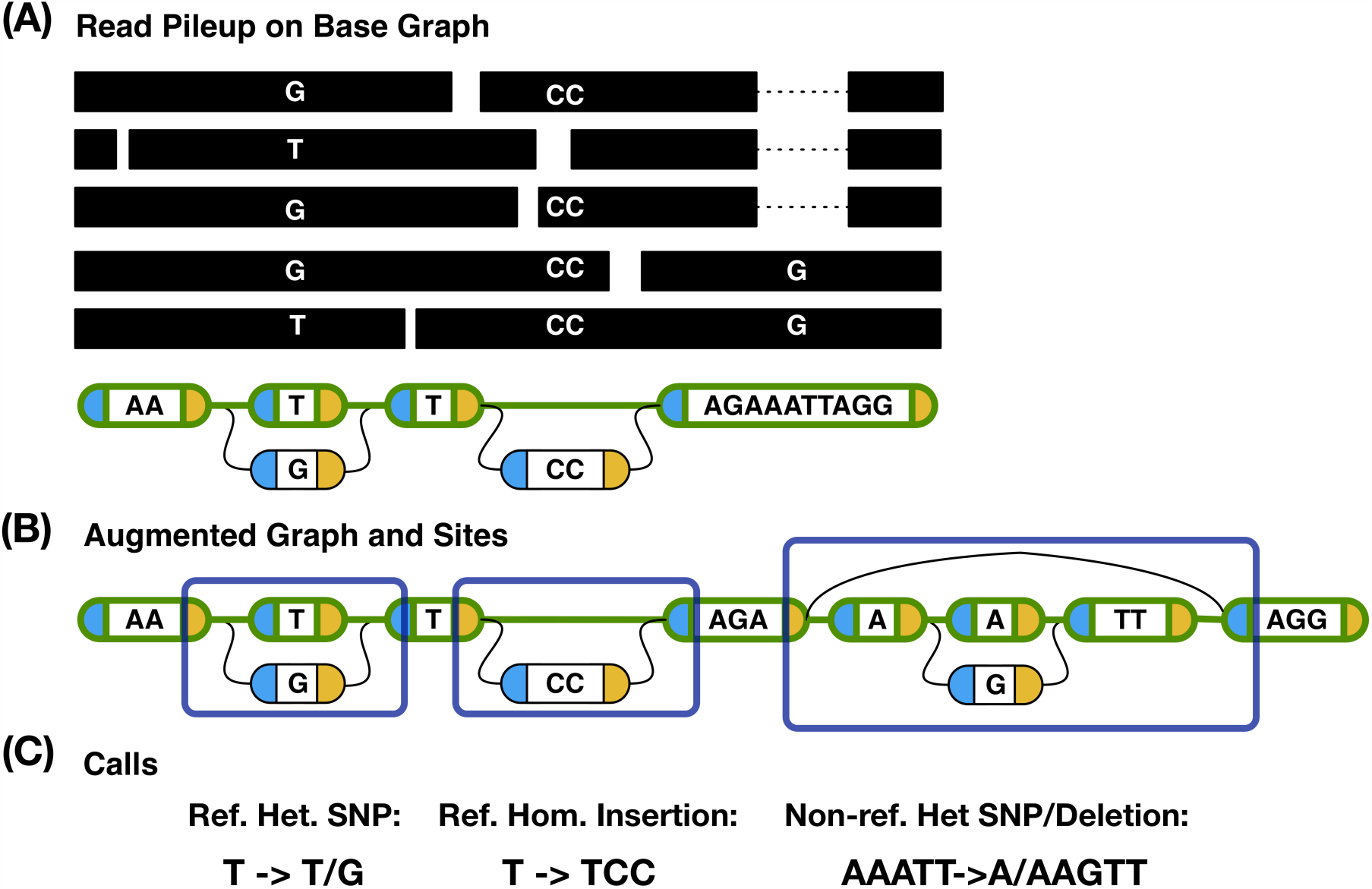
Variant Calling with Genome Graphs. (A) Read pileup on a base graph whose reference path is highlighted in green. Only variant or non-reference base values are shown in the reads. (B) The augmented graph contains the base graph as well as new structures implied by the pileup. This graph contains three top-level ultrabubbles, each forming a site. (C) Variant calls for each site. The first two (a heterozygous SNP and a homozygous insertion) are considered reference calls because they were present in the base graph, whereas the third (a heterozygous combination of a SNP and a deletion) is non-reference because it was novel to the augmented graph.

We compared the results from the graph variant calling pipeline with the Platinum Genomes benchmark data for samples NA12877 and NA12878^32^ using vcfeval, which corrects for VCF representation ambiguity by comparing at the haplotype level ^33,34^. To provide additional controls, Freebayes ^35^, Platypus ^36^ and Samtools ^30^ were run on BWA-MEM ^6^ alignments of the same input data to GRCh38 with their default options in order to produce positive control variant calls. Figure 4 (A) shows the precision and recall of each method aggregated across both samples and all regions. Figure 4 (C) and (D) show precision-recall curves for SNPs and indels, respectively. In comparison to the primary graph (the graph containing only the existing reference sequence, and therefore a control for the same variant calling algorithm applied to just the knowledge in the existing reference), the 1KG and Cactus graphs’ F1-scores (Supplementary Table 1) increased by 3.50% and 1.98%, respectively, increasing for both single nucleotide variants (3.13%, 1.95% respectively) and indels (6.02%, 4.40% respectively). Furthermore, 1KG graphs have the overall highest accuracy (F1 score) of all methods, although Samtools and Platypus perform best overall for SNPs and indels, respectively. Supplementary Section 7 contains additional breakdowns of the F1-scores by region (Supplementary Figs. 7-8), sample (Supplementary Fig. 9), and type (Supplementary Fig. 10), as well as scores without clipping to confident regions (Supplementary Fig. 11). Generally (in 13 out of 18 cases), the 1KG graph has higher accuracy than both the primary and scrambled controls.

**Figure 4:**
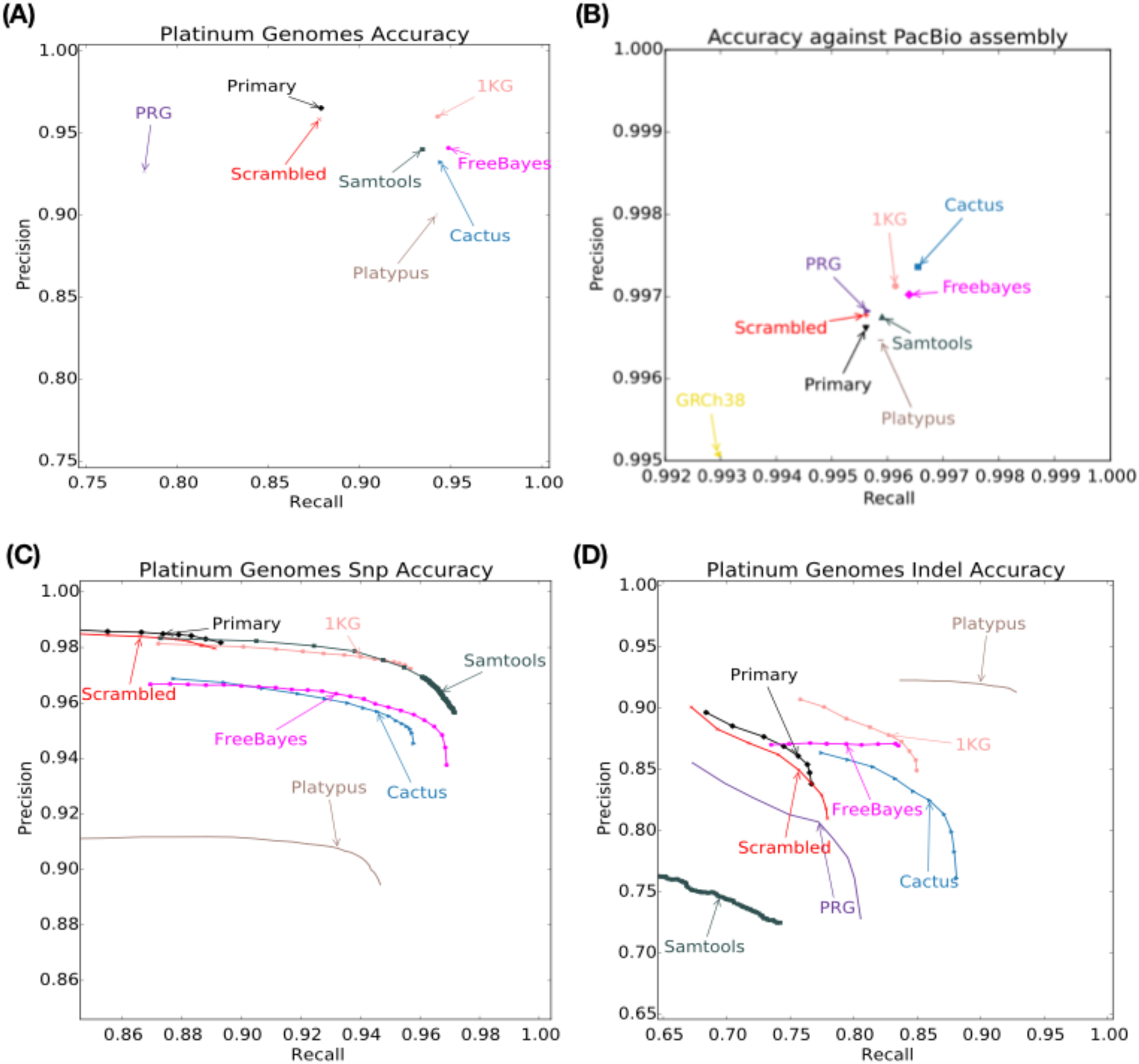
Variant Calling Evaluation. (A) Precision (portion of called variation in agreement with the truth set) and recall (portion of variation in the truth set in agreement with what was called) against the Platinum Genomes truth VCFs aggregated across NA12877 and NA12878 for all regions, as measured by vcfeval. (B) Per-base precision and recall as measured by the reference-free evaluation in BRCA1, BRCA2, LRC_KIR, and MHC. The GRCh38 point shows a comparison of the existing primary reference haplotype sequence against the de novo assembly. (C) - (D) Breakdown of precision and recall from (A) into SNPs and indels, respectively. Curves are shown by including accuracies at quality thresholds that fall within a radius of 0.1 around the maximum F1. Full results featuring F1-scores for all graphs are in Supplementary Section 7.

We define a *reference call* as a call asserting the presence of a position or edge in the base graph. The experimental graphs can dramatically reduce the number of non-reference calls, as compared to control. For example, the Cactus and 1KG graphs reduce non-reference calls by more than a factor of ten (Fig. 5 (A)) relative to the Primary reference graph. Furthermore, the precision of these reference calls is higher than the non-reference calls for the non-scrambled graphs (Fig. 5 (B)).

**Figure 5:**
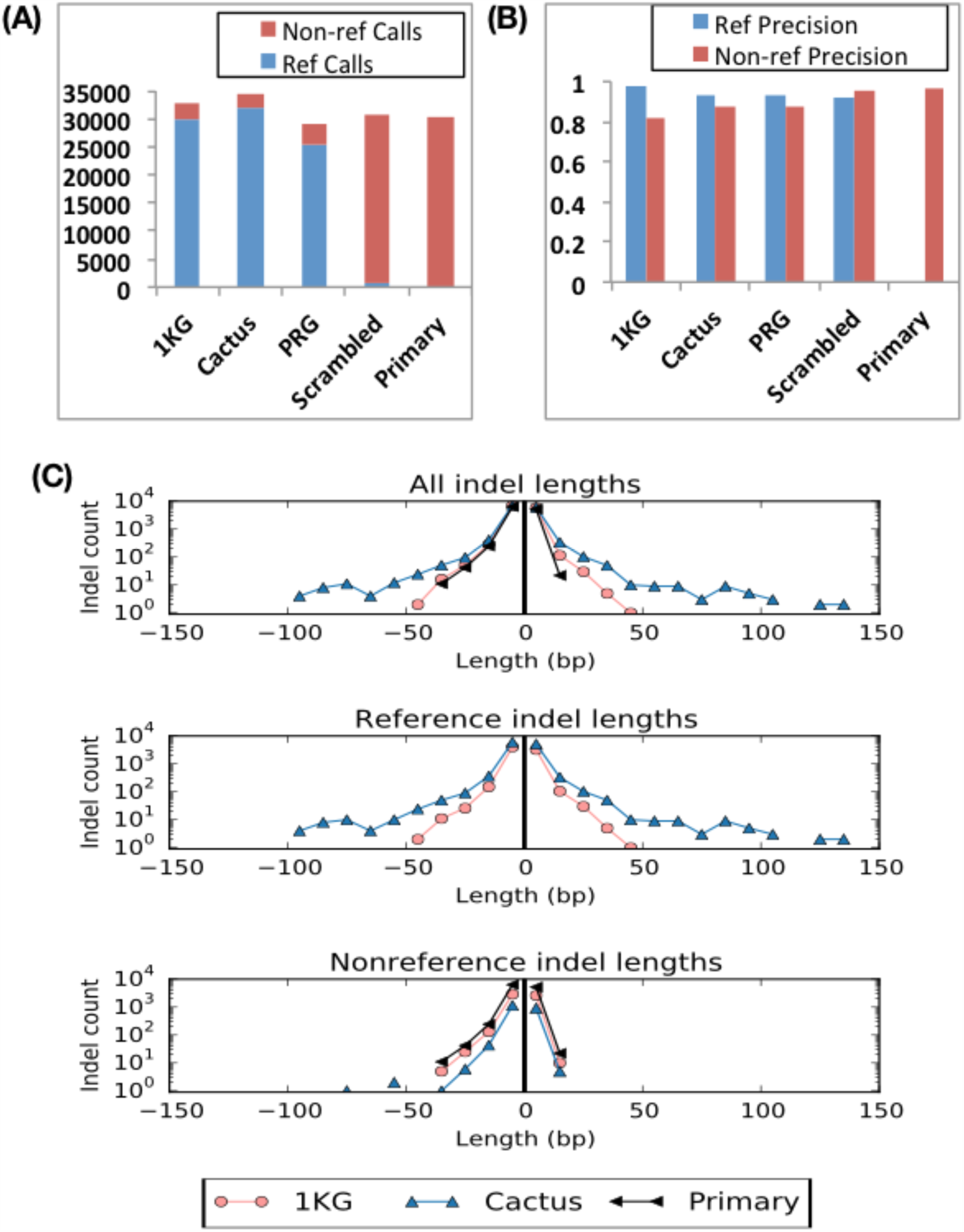
Reference versus Non-reference Calls. (A) Total number of reference and non-reference calls across all samples and regions. (B) Precision of reference and non-reference calls. (C) Indel lengths of reference and non-reference calls, where insertions and deletions are represented by positive and negative lengths, respectively. In all cases we ignore calls of GRCh38 reference alleles, as these numbers are reported from GRCh38-based output VCFs.

Larger structural variants can be called using the same logic as point mutations, provided they are already in the graph; Figure 5 (C) displays the indel length distribution for the two top-performing graphs and the primary control, as well as a breakdown of indel lengths for reference and non-reference calls. The reference call indel lengths in the experimental graphs are larger than the Primary and non-reference lengths and, in the case of Cactus, contain indels exceeding the read length. This adds up to a large number of additional called bases: for example, across the regions the Cactus graphs call 94 indel events larger than 50 base pairs totaling 10045 bases, none of which are found using the Primary graph with the same algorithm.

To mitigate potential biases with the Platinum Genomes benchmark data as a truth set^32^, we conducted what we term a “reference-free” evaluation of vg’s variant calling accuracy, by comparing against de novo assemblies instead of assumed-true variant calls. In brief, short reads pooled from two haploid assemblies were used to call variants on each sequence graph. The accuracy of this reconstruction was evaluated using PacBio-based *de novo* assembly fragments, which by definition are free of reference artifacts and are derived from a different sequencing technology (see Online Methods, Supplementary Section 8 and Supplementary Figure 12). The results can be seen in Figure 4 (B) and Supplementary Figure 13. Several experimental graphs have greater precision and recall than the Scrambled and Primary controls; combined across all regions except SMA (which was insufficiently covered by PacBio assemblies to be usefully analyzed), vg on the Cactus graph outperformed existing methods. The results appear to agree closely with those from the VCF-comparison-based evaluation, considering that the two techniques use different sources of truth and different evaluation metrics.

## Short Path Accuracy

We sought to understand how complete and accurate the sequence graphs studied were in their representation of common variants. To approximate this, we measured the fraction of lightly error-pruned K-mer instances (here K=20, see Online Methods) in a large subset of 1KG sequencing reads that were present within each graph, calling this metric *K-mer recall* (see Online Methods). We observe (Fig. 6, Supplementary Section 9, and Supplementary Fig. 14) that graphs built from the largest libraries of variation contain the great majority of such K-mer instances. For example the 1KG, PRG and Cactus graphs contain an average across regions of 99% of K-mer instances, while the primary graph contains an average across regions of 97%. Conversely, we asked what fraction of 20-mer instances present in a graph were not present in any 1KG read, calling this metric *K-mer precision*. Strikingly, we find that precision is greatly reduced in some graphs relative to control. For example around 15% (averaged across regions) of 20-mers enumerated from 1KG graphs do not appear in any 1000 Genomes low coverage read. We hypothesize that this is because the density of variation is sufficient in such graphs to admit many paths implying recombinations that are either absent or very rare in the population. To attempt to raise precision, for the 1KG data we constructed graphs using haplotype information to eliminate a substantial subset of unobserved paths, creating the “1KG Haplo 50” graph (Supplementary Section 10). This resulted in increased precision (by about 10 percentage points in BRCA2) with only a small reduction in recall, as shown in Figure 6 and Supplementary Figure 13. However, this comes at the cost of a performance degradation in read mapping (Fig. 2) and variant calling (Supplementary Section 7). One possible explanation for the performance reduction is that the necessary duplication (“unmerging”) of paths in this procedure reduced the aligner’s ability to unambiguously map reads.

**Figure 6:**
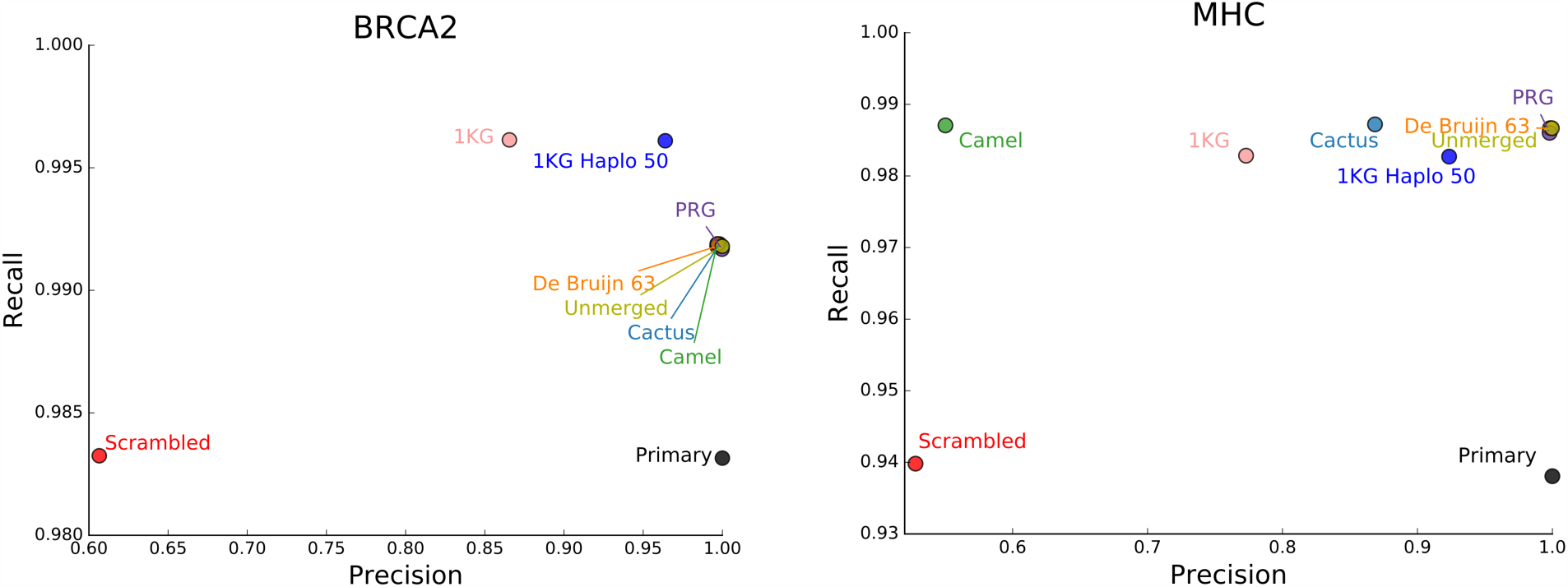
Short path completeness and accuracy. Assayed by comparing 20-mer instances.

## Graph Character

We found that even within each region the different submitted graphs varied substantially in their performance on our evaluations of read mapping and variant calling. They varied even more so with respect to basic graph properties (Supplementary Section 11, Supplementary Tables 2-9). To quantify this variability we defined normalized graph metrics for basic graph properties. *Graph compression* is the length of the primary reference sequence within the region divided by the sum of the lengths of the nodes in the graph. It is a normalized measure of the number of positions in the graph. The (*base*) *degree* is the average per-side degree of the graph in a bidirected graph representation with single-base nodes, and is a measure of how much branching a graph contains. The *cut width* (Supplementary Table 10) is a measure of apparent sequence rearrangement. Briefly, within a topologically sorted graph, where all positions are ordered, cut width is the average over all gaps between successive positions of the number of edges connecting positions on the left side of the gap to positions on the right side of the gap (Supplementary Section 12)^37^. We see wide variation in these measures across the graphs (Fig. 7). Furthermore, across the different regions we find that there is an inverse correlation (R=-0. 674, p=0.00230) between cut width and variant calling accuracy and a positive correlation (R=0.244, p=0.0268) between compression and variant calling accuracy (Supplementary Fig. 15). The base degree does not significantly correlate with variant calling accuracy. These correlations suggest that uncompressed and highly rearranged graphs do not work effectively with our current read mapping and variant calling process.

**Figure 7:**
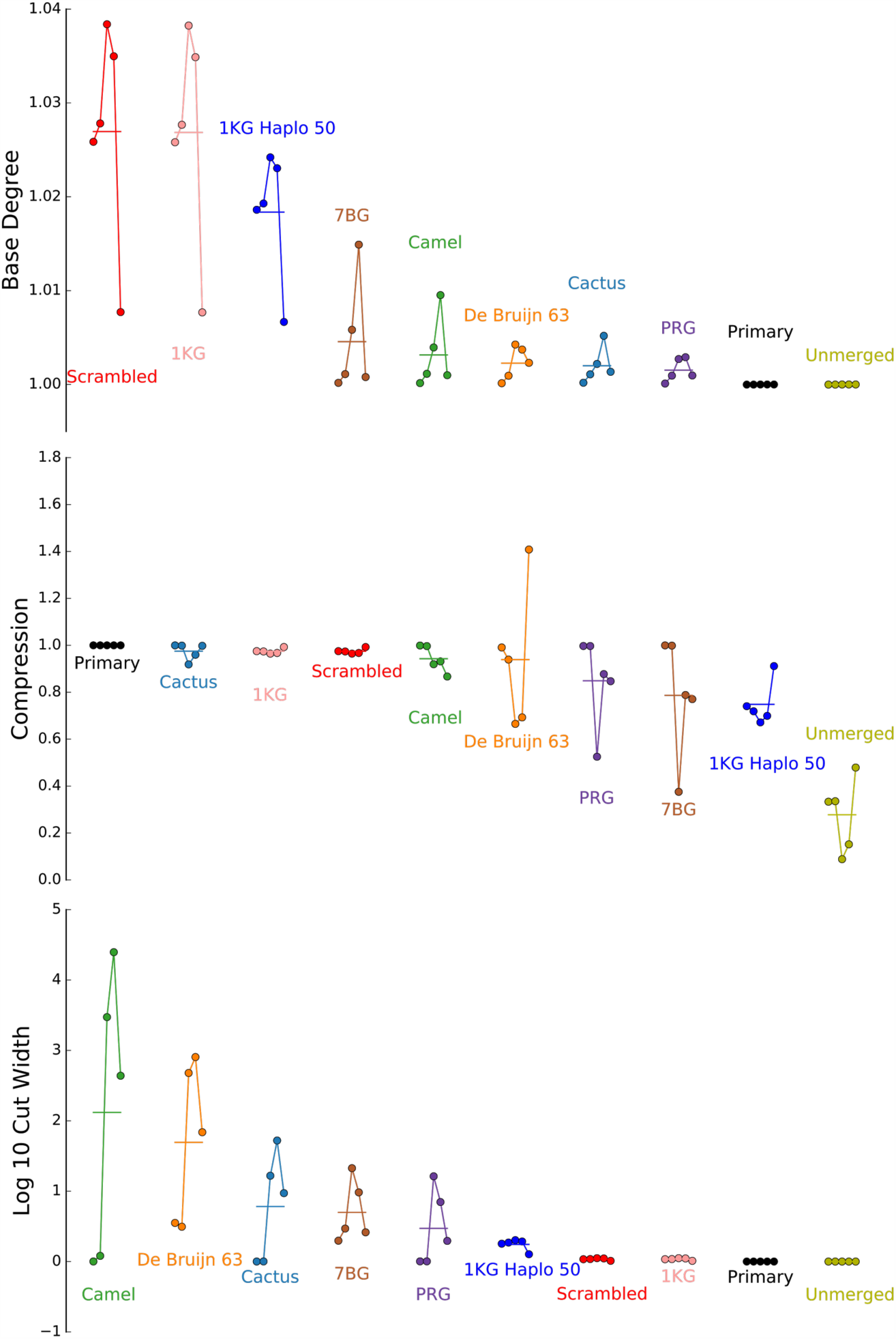
Empirical graph statistics. In each panel the result for each region is shown by a dot, in the following order: BRCA1, BRCA2, LRC_KIR, MHC, and SMA.

## Discussion

Contemporary non-graphical variant calling procedures use different algorithms for each class of variants: substitutions, small indels, larger indels, balanced rearrangements, and so on. We have demonstrated that variant calling on a sequence graph mostly obviates this complexity, because being able to call the presence or absence of elements within a sample graph is potentially sufficient for calling known structural and point variation equally well. The simple, nascent variant calling algorithm we tested produced variant calls that were quite concordant with those from other state-of-the-art variant calling pipelines, while unifying the calling of known SNPs and other known structural variation. That individual tools slightly outperformed the variant calling algorithm presented here in terms of individual variant types, i.e. snps and indels, is unsurprising given the relative maturity and algorithmic sophistication of those tools. Importantly, many of the submitted graphs showed improved variant calling performance over the primary and scrambled graphs. The relative improvements come alongside a large reduction in the number of non-reference calls. Furthermore, reference calls were more accurate than non-reference calls, suggesting that variant calling is indeed more accurate overall when the variants themselves are contained within the graph. These results support the notion that sequence graphs can transform variant calling by reducing it to the simpler problem in which only rare variants, absent from the graph, must be discovered de novo. It is possible to foresee cutting the number of non-reference point variation calls from the millions, as in standard genome wide pipelines today, to on the order of thousands (see Supplementary Section 3).

During the course of the variant calling comparison, we developed an appreciation for the shortcomings of relying solely on the Platinum Genomes benchmark data as a truth set^32^. A key concern is that the Platinum Genomes calls were derived by means of a consensus of contemporary methods, all of which use the existing linear reference and BWA-MEM-based mappings. Additionally, compared to vg, the Platinum Genomes dataset often uses different combinations of calls to “spell” the same haplotype. Moreover, it often omits calls necessary to spell a haplotype because it is not confident in them. While the omitted calls are in regions marked as low confidence, a variant normalizer cannot normalize a call that is not there. To get around these problems and potential biases we introduced a reference-free method for assessing variation calls. This evaluation demonstrated good consistency with the Platinum Genomes in terms of the relative ranking of the different methods evaluated, and demonstrated clearly that the best graph methods slightly outperform existing methods.

Supporting the observed improvements in variant calling, we demonstrate that read mapping can be made both more comprehensive and less ambiguous with sequence graphs. Increases in perfect mapping and reductions in substitution and indel rates were broadly consistent with the effect we would expect if the graphs were representing the majority of common polymorphisms, leaving the residual read error rates to account for the majority of alignment differences. In this sense read mappings were demonstrated to be less locally ambiguous, with mismatches and edits having a more clearly defined meaning. Furthermore, the fact that read mappings were also less globally ambiguous (i.e. more certain in their overall placement within the genome) is perhaps surprising. We thought at the outset that using detailed graphs would have the drawback of increasing the number of times a read maps to two or more places by increasing the sheer number of mapping possibilities. However, we found that the opposite is true - the addition of known polymorphisms to the graph allows reads to better distinguish their true mapping location from secondary, paralogous locations. Scaled genome-wide, these improvements could help canonicalize mapping to the vast majority of variation, which will become especially important as genome variants are increasingly used in the clinic. The increases in perfect mapping could also allow alignment to be made more efficient by allowing larger, more stringent seeds or more aggressive ungapped matching. Our early work with vg indicates that there is ample opportunity for improvement and investigation of these novel approaches to the design of high-performance mapping algorithms. We also collected some preliminary data that suggests that the gains in mapping obtained by moving from the existing reference to a graph like the 1KG graph are super-population specific, suggesting that sequence graphs have the potential to reduce the local ethnic bias inherent in a single reference genome.

By taking a community approach, we were able to sample a wide variety of the contemporary software for building sequence graphs. It is apparent that different methods produce dramatically different graphs, as measured both by direct graph analysis and by practical performance as a reference for common genomics tasks, suggesting that the field is just in its formative stages. In trying to understand how “complete” and “accurate” graphs built with today’s methods are at representing short sequences present in the population, we encountered several surprises. In particular, we found a large number of short non-biological paths created within the highest degree graphs, such as the de Bruijn graphs, parts of the 1KG graphs, and certain of the Seven Bridges graphs. We tried modifying the 1KG graphs to reduce the number of false recombination possibilities without much success. We may in the future find that we can tolerate these short non-biological paths, or that another approach is needed to eliminate them.

One alternative approach is to create uncompressed, lower-degree graphs by duplicating variable regions to directly represent haplotypes, but it is likely that, as demonstrated by the 1KG Haplo 50 and (at the logical extreme) Unmerged graphs, the resulting long, equivalent sequence paths would create too much multi-mapping ambiguity. Perhaps a better solution may be the use of haplotype information embedded within the sequence graph^38^, making it a *variation graph*. This would allow algorithms to map to a common graph coordinate system while accounting for variants, read errors, and recombinations within the mapping process itself. This approach would eliminate the need for several inelegant heuristics used in contemporary linear-reference-based analysis pipelines ^28,39^.

Sequence graphs can now be built from libraries of common variants, and tools like vg, though still experimental, illustrate the huge potential of the graph-based approach. There are a number of questions yet to be tackled. How should duplications and repeats be represented? How can one best map to a graph? How should short variants whose homologies are unclear be parsed? How can graphs be used to enable a more comprehensive taxonomy of variation? These questions all represent open avenues for future research.

## Online Methods

### Source Data

Participants were provided with a data set consisting of five genomic regions (BRCA1, BRCA2, LRC_KIR, SMA, and MHC) to use in the creation of their graphs. The data set came in the form of a “reference” sequence and one or more “alternate” sequences for each region. For the LRC_KIR, SMA, and MHC regions, those alternate sequences were the alt loci present in GRCh38.p2 for the regions of the same names in the assembly, with the reference being the portion of the corresponding chromosome encompassing the chromosomal coordinates for all of the alts. The reference regions for BRCA1 (ID 672) and BRCA2 (ID 675) were downloaded from Entrez Direct, while alternate sequences were the annotated genes from the CHM1 hyatidiform mole assembly, and the LRG sequences for those genes ^40–42^. Some participants used additional source data in constructing their graphs.

### Graph Format

All graphs were generated in or converted into an SQL text format for submission. The graphs were then loaded into databases compatible with the GA4GH Graph Reference Server, and servers for the graphs were hosted on a Microsoft Azure cloud instance. Individual evaluation tools hit against these API endpoints. For read alignment and variant calling purposes, graphs were downloaded from the servers using the sg2vg converter tool, written for this project, and stored in .vg graph format. This on-disk format could be efficiently indexed for read alignment—a function that the GA4GH server did not support—and so was preferred for evaluations dependent on read alignment. The graphs themselves were created using a variety of methodologies and approaches, detailed in Supplementary Section 10.

### Alignment Target Quality

The submitted graphs were used to align reads from 2,691 low-coverage samples from the 1000 Genomes project, which had been aligned to GRCh38 with BWA-MEM ^6^. Alignments to the primary reference and, where available, the GRCh38 alt loci for a region were downloaded using Samtools ^30^. The process took advantage of the tool’s ability to subset indexed files over FTP in order to obtain just reads mapped within the region ^30^. Next, the alignments were converted into reads, yielding the relevant reads for that sample and region. Unpaired reads in the downloaded set were discarded. An attempt was made to correct for a known data corruption bug in the version of BWA-MEM used to produce the alignments, by taking the sequences given for alignments to the primary reference over the sequences given for the same read aligned to an alt, where available (Heng Li, personal communication). Input graphs were downloaded from the reference servers using the sg2vg program. They were then broken into nodes of no more than 100 bases each and re-numbered according to a heuristic extension of topological sort to cyclic graphs. Graphs were indexed and alignment was performed with the vg program, using a K-mer size of 16 and an edge-crossing limit of 3 for the GCSA2 index. The portion of reads mapping uniquely was calculated. To qualify as uniquely mapped, a read had to have a primary mapping with 0.95 matches per alignment column or fewer. Additionally, qualifying reads had to have either no secondary mapping or a secondary mapping having fewer than 0.85 matches per column. The denominator for the portion mapping uniquely was the number of reads having either a secondary mapping distinct from the read’s primary mapping or no secondary mapping at all (see Supplementary Section 3). The portion of reads mapping perfectly was defined as the portion having 1 match per alignment column. The substitution rate was defined as the portion of bases in length-conserving replacements out of all substituted or matched bases. Bases matched or substituted against N characters in the reference graph were ignored. The indel rate was defined as indel count divided by substituted and matched bases. Bases matched or substituted against reference Ns were ignored, as were indels that constituted softclips.

### Platinum Genomes Variant Calling Evaluation

A graph variant calling pipeline based on the Samtools pileup method was implemented in vg and run independently on three 50x coverage samples from Platinum Genomes (NA12877-9). First, the reads were mapped to each graph as described above. The alignments were then filtered to remove secondary mappings, as well as mappings with mapping quality score less than 15, mappings that had been promoted to primary over another properly paired mapping of greater single-end score, and mappings with soft-clipped or ambiguous ends (more details in Supplementary Section 7). A pileup of aligned read bases was then constructed for each position and edge in the graph ignoring bases with read quality score less than 10. The SNPs, insertions, and deletions implied by the two most supported non-reference entries in each pileup were then added into the graph to create an “augmented” graph. Sites in the augmented graph were computed using the ultrabubbles algorithm^31^. For each site, the two non-reference paths with the most read support were greedily chosen using a breadth-first search. A path’s read support was defined here as the minimum pileup support of any node or edge it contains; each node’s support was calculated as the average support across the node’s bases. Finally, given the reference path and these two alternate paths for each site, a genotype was computed using a thresholding heuristic based on the ratios of the paths’ pileup supports. Alternate alleles were called as heterozygous if they had at least three times as much read support as the reference (or six times for a homozygous alt call). The genotypes were written directly to VCF. The variants were normalized by using vt^43^ to flatten multibase alts that contain reference calls. Calls for both NA12877 and NA12878 were compared against their respective Platinum Genomes truth set VCFs; these were the only samples with truth VCFs available. Precision and recall against the truth set were assessed with vcfeval^33^. True and false positives and negatives returned from this tool were classified as SNPs and indels using bcftools, and clipped into the Platinum Genomes confident regions. Precision, recall and F1-scores were then computed for each possible quality threshold in the VCF. For the vg call results, minimum read support (AD field in VCF genotype) across called alleles was used as a proxy for quality. Aggregate results across samples and regions were computed by pooling the vcfeval results together. The precision-recall curves (Fig. 4 (C) and (D)) were drawn by filtering the VCF files by all values of variant quality and displaying only those within distance 0.1 of the maximum F1-score. The points shown in Figure 4 (A) were chosen to correspond to the quality threshold yielding the maximum F1-score.

### Reference-Free Evaluation

A “synthetic diploid” genome was conceptualized by combining data from two haploid samples, CHM1 and CHM13^44^. For each sample, GRCh38-aligned low-coverage Illumina reads and relatively complete PacBio-derived assemblies were obtained. The CHM1 and CHM13 reads were obtained by combining both runs from NCBI SRA SRX1391727 and SRX1082031, respectively, and mapping to GRCh38 using BWA-MEM^6^. The CHM1 assembly used was GenBank accession number GCA_001297185.1, while the CHM13 assembly was GCA_000983455.2. For each region, a pooled collection of the relevant Illumina reads across both CHM1 and CHM13 was created. Next, the reads were subsampled for balanced coverage between the two haploid genomes as would be expected in a real diploid sample. For each submitted graph under tests, the reads were aligned using vg, and the vg variant caller was used to produce variant calls. The resulting VCF for each graph construction method and region combination was turned into a new “sample graph” to which the relevant portions of the PacBio assemblies were aligned. Treating the aligned assembly fragments as the truth, the precision and recall of each sample graph were measured as a function of which original submitted graph it was derived from.

Assembly fragments used for evaluation were selected by alignment of the primary reference sequences for the regions against the CHM1 and CHM13 assemblies using BLAT version 36x2 ^45^. Aligned regions in the assembly covering more than either 50% of an assembly contig or 50% of a region, with more than 98% identity, were extracted from the assembly and used for realignment. The SMA region was excluded from the evaluation due to patchy, overlapping coverage of the region in the two assemblies. Additionally, the first 87,796 bases of the LRC_KIR region were excluded from the sample graphs and the aligned truth set contigs due to an apparent lack of representation in the CHM13 assembly.

### Assessing Graph Completeness

Reads aligning to the test regions were obtained from 2,691 low-coverage samples in the 1000 Genomes Project, and each sample’s reads were used to generate a collection of K-mers (K=20) using Jellyfish ^5,46^. These were compared against the collection of K-mers in each graph as enumerated by vg with an edge-crossing limit of 7. In order to account for K-mer frequencies, duplicate K-mers were *not* ignored. K-mers containing N characters were ignored in both collections, and K-mers only observed once in their sample were ignored in the 1000 Genomes-derived K-mer collection. This latter filter was intended to remove the large majority of erroneous K-mers: we expect errors to be Poisson-distributed one-off events while real variants are likely to recur within a sample. Recall, defined as the portion of all read-derived K-mers present among the graph-derived K-mers, and precision, defined as the converse, were computed for each graph.

### URLs

VG, https://github.com/vgteam/vg.

Patches to VG, https://github.com/adamnovak/vg/tree/graph-bakeoff.

GA4GH to VG Importer, https://github.com/glennhickey/sg2vg.

VG to GA4GH Exporter, https://github.com/glennhickey/vg2sg.

GA4GH Graph Schemas, https://github.com/ga4gh/schemas/tree/refVar-graph-unary.

GA4GH Graph Server, https://github.com/ga4gh/server/tree/graph.

Graph evaluation software, https://github.com/BD2KGenomics/hgvm-graph-bakeoff-evaluations.

FASTG, http://fastg.sourceforge.net/.

Illumina Platinum Genomes, http://www.illumina.com/platinumgenomes/.

Jellyfish, http://www.cbcb.umd.edu/software/jellyfish/.

Platypus, http://www.well.ox.ac.uk/platypus.

Freebayes, https://github.com/ekg/freebayes.

Samtools, http://www.htslib.org/.

VCFeval, https://github.com/RealTimeGenomics/rtg-tools.

### Software Versions and Commit Hashes

VG, 158d542497445b532b0e9e40223f5023ee6b52dd.

GA4GH to VG Importer, 468026ad70f0425af1959b287ffcaac91b8a9deb.

VG to GA4GH Exporter, 4efde8e64a8bd113a0e83685628bbaf0cbc2be3f.

GA4GH Graph Schemas, ea58ac46dad84be67c500e517ff2fb05a43a187a.

GA4GH Graph Server, c6daebca4c69a4ff4d9d56cfdf587556f2ce1116.

Graph evaluation software, 52b0537713629471f6ea97ccf552d6727c630f3d.

Freebayes, 9e983667d47f6b5dcbb90070da8de69714038f46.

Samtools, version 1.3.1.

## Acknowledgments

This work would not have been possible without the generous support of the National Human Genome Research Institute (1U54HG007990 [BD2K] to B.P. and D.H., 5U41HG007234 [GENCODE] to B.P.); the W. M. Keck Foundation (DT06172015 to B.P. and D.H.); the Wellcome Trust (100956/Z/13/Z to G.M.); the Simons Foundation (SFLIFE# 351901 to B.P. and D.H.); the ARCS Foundation (2014-15 ARCS fellowship to A.M.N.) and Edward Schulak (Edward Schulak Fellowship in Genomics to A.M.N.).

## Author Contributions

A.M.N. contributed the Camel graphs, wrote read mapping and variant calling code for vg, ran the read mapping and reference-free evaluations, and contributed extensively to the organization of the manuscript. G.H. contributed the Cactus, 1KG, 1KG Haplo 50, Primary, and Scrambled graphs, and performed the variant calling evaluation. E.G. contributed the bulk of the vg tool. S.B. contributed the analysis of graph statistics. A.C., S.G., N.O., and A.W.Z. contributed the Curoverse graphs, with A.W.Z. supervising. A.D., J.K., supervised by G.MV., contributed the PRG graphs. J.E. contributed alignment code to the vg tool. M.A.S.E. contributed advice and corrections to the manuscript. A.K. contributed the De Bruijn 63 graphs. S.K. contributed code to the GA4GH graph server and contributed extensive organizational support. D.K. and G.R. contributed the SBG graphs. H.L. contributed experimental design support for the read mapping evaluation, and advice on the manuscript. M.L. worked on scaling up the variant calling pipeline to whole genomes. K.M. contributed a set of graphs for the chromosome X centromere. M.S-O. contributed code to the GA4GH graph server and managed the graph data import pipeline for this work. R.D. contributed to the design of the GA4GH graph server interface and schemas, and supervised other authors. G.M., D.H., R.D. and B.P. contributed to the design of the project and supervised authors. B.P. and organized the project directly, and wrote extensive portions of the manuscript. All authors edited the manuscript.

